# Timescales of spontaneous fMRI fluctuations relate to structural connectivity in the brain

**DOI:** 10.1101/655050

**Authors:** John Fallon, Phil Ward, Linden Parkes, Stuart Oldham, Aurina Arnatkevic̆iūtė, Alex Fornito, Ben D. Fulcher

## Abstract

Intrinsic timescales of activity fluctuations vary hierarchically across the brain. This variation reflects a broad gradient of functional specialization in information storage and processing, with integrative association areas displaying slower timescales that are thought to reflect longer temporal processing windows. The organization of timescales is associated with cognitive function, distinctive between individuals, and disrupted in disease, but we do not yet understand how the temporal properties of activity dynamics are shaped by the brain’s underlying structural-connectivity network. Using resting-state fMRI and diffusion MRI data from 100 healthy individuals from the Human Connectome Project, here we show that the timescale of resting-state fMRI dynamics increases with structural-connectivity strength, matching recent results in the mouse brain. Our results hold at the level of individuals, are robust to parcellation schemes, and are conserved across a range of different timescale-related statistics. We establish a comprehensive BOLD dynamical signature of structural connectivity strength by comparing over 6000 time-series features, highlighting a range of new temporal features for characterizing BOLD dynamics, including measures of stationarity and symbolic motif frequencies. Our findings indicate a conserved property of mouse and human brain organization in which a brain region’s spontaneous activity fluctuations are closely related to their surrounding structural scaffold.

## 1 Introduction

The brain’s complex spatiotemporal dynamics unfold on an intricate web of axonal connections: the connectome [1, 2]. These pathways facilitate information transfer between brain regions, manifesting in a complex relationship between connectome structure and neural dynamics. Reflecting the pairwise (region–region) nature of structural connectivity, existing studies have overwhelmingly compared pairwise measurements of anatomical connectivity to pairwise statistical relationships between neural activity time series, or functional connectivity, often using simulations of network dynamics to better understand how the observed relationships may arise [3–19].

Structural connectivity is highly informative of functional connectivity, consistent with the connectome as a physical substrate constraining inter-regional communication dynamics. However, our understanding of pairwise structure–function relationships remains disconnected from our understanding of how a brain area’s structural connectivity properties shape its local activity dynamics. The structural connectivity profile of a region’s incoming and outgoing axonal connections characterizes its function [20]. Furthermore, the activity dynamics of brain areas follow a functional hierarchy, with rapid dynamics in ‘lower’ sensory regions and slower fluctuations in ‘higher’ regions associated with integrative processes [21–25]. The spatial variation of intrinsic timescales has been measured using ECOG [22], MEG [26–28], TMS–EEG [29], and fMRI [25, 30–35], and may functionally correspond to a variation in temporal receptive windows [timescales over which new information can be actively integrated with recently received information [21, 35, 36]]. Spatial variation in intrinsic activity fluctuations may form a key basis for the brain’s functional hierarchical organization, shaped by variation in the brain’s microcircuitry [37–39]. This organization is thought to be important for behavior and cognition [21, 40–42] and its disruption has clinical implications; e.g., differences in intrinsic timescales are associated with symptom severity in autism [35]. While much is known about the structure– function relationship at the level of pairs of brain regions, and how structural and functional connectivity architecture shape cognitive function and are affected in disease [43–46], relatively little is known about how structural connectivity affects the information-processing dynamics of individual brain areas. In particular, we do not yet understand the role structural connections play in the organization of timescales.

Recent work has provided statistical evidence for a relationship between regional tract-tracing estimates of anatomical connectivity and rs-fMRI dynamics in the mouse brain [47]. This work took a comprehensive, data-driven approach, comparing over 7000 properties of regional BOLD dynamics (using the *hctsa* software package [48, 49]) to three key structural-connectivity properties—degree, betweenness, and clustering coefficient—measured in each of 184 brain areas. The tract-traced connectivity measurement available in mouse [50] also allowed the role of directed and weighted connectivity information to be investigated. The weighted in-degree, 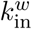, showed the strongest correlation to BOLD dynamics, particularly with its auto-correlation properties (including the Fourier spectral power in different frequency bands). For example, relative high-frequency power (*f* > 0.4 Hz) was found to be negatively correlated to weighted in-degree, 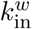) (*ρ*_*V*_ = −0.43, partial Spearman correlation controlling for region volume). The results suggest that structural connectivity may play a role in the spatial patterning of intrinsic timescales: brain areas with a greater aggregate strength of axonal input (highest 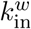) display slower timescales of spontaneous activity fluctuations, consistent with the predictions of model simulations [32, 51, 52]. Despite the low sampling rate of rs-fMRI, recent work has shown a strong correlation between timescales estimated from EEG and fMRI [35], suggesting that a similar trend may hold at much faster timescales. When ignoring edge directionality, Sethi et al. [47] found weaker but statistically significant relationships between (undirected) weighted-degree, *k*^*w*^, and rs-fMRI dynamics. This suggests that a similar relationship may hold in human, where the directionality of connections cannot be measured through non-invasive MRI methods like diffusion-weighted imaging.

We know of only two investigations into how structural connectivity properties relate to BOLD dynamics in the human cortex, and both have reported weak relationships between structural-connectivity strength and: (i) low-frequency rs-fMRI fluctuations, Pearson’s *r* = 0.12 [34]; and (ii) the log-linear slope of the Fourier power spectrum, *r* = 0.22 [31]. These weak correlations may be due to both studies measuring BOLD for a short duration (less than 300 volumes at a sampling rate TR = 2.5 s) in a small sample of individuals: 30 [31] and 36 [34]. However, neither study controlled for region volume, which correlates strongly with the autocorrelation properties of the BOLD signal, due to the averaging of more voxel-wise signals in larger areas [47, 53]. The Human Connectome Project (HCP) dataset alleviates many of these issues, containing a large rs-fMRI dataset collected at a high sampling rate, TR = 0.72 s, across 1200 time points [54].

Here we characterize the rsfMRI signature of structural connectivity in the human cortex using data from the HCP. We aimed to investigate whether more strongly connected regions are associated with slower timescales of BOLD activity in the human cortex, as they are in the mouse brain [47]. Following the results in mouse, we use relative low-frequency power (RLFP) to estimate the prominence of slow BOLD fluctuations (*f* < 0.14 Hz), and show that this measure is strongly correlated with other measures of timescale obtained from the decay of the autocorrelation function with time-lag [24, 35]. Consistent with predictions from mouse, RLFP increases with structural connectivity strength in human cortex, *ρ*_*V*_ = 0.53 (partial Spearman correlation adjusting for region volume, *p* = 2×10^−3^). Our results hold at both the group and individual level, and across different cortical parcellations, reflecting a robust interspecies conservation of how structural connectivity and regional activity dynamics are related.

## 2 Results

### 2.1 Methods Summary

We investigated whether a brain area’s structural connectivity strength is related to its spontaneous BOLD dynamics, as illustrated schematically in Fig. 1. Our methods are summarized briefly here (and detailed in Methods). We used an HCP dataset of 100 healthy, unrelated participants (54 male, 46 female; 22– 35 years old) [54]. Our main analysis focuses on the left hemisphere of the 68-region Desikan-Killiany Atlas [55] (analysis of the right hemisphere yielded similar results, see below). Structural connectivity was estimated from the diffusion data using `MRtrix3` [56] and the FMRIB Software Library [57], performing tractography with 10 million streamlines using FACT, ACT, and SIFT-2, yielding a 34 × 34 left-hemisphere connectome. Following work in mouse [47], we summarized the structural connectivity of each brain area as its node strength, *s*, estimated as the total number of diffusion MRI-reconstructed streamlines attached to it (equivalent to weighted degree, *k*^*w*^). rs-fMRI data were processed after regressing standard nuisance signals (including the global signal) and were high-pass filtered at 8 × 10^−3^ Hz, yielding a 34 × 1200 (region × time) fMRI data matrix.

**Figure 1:**
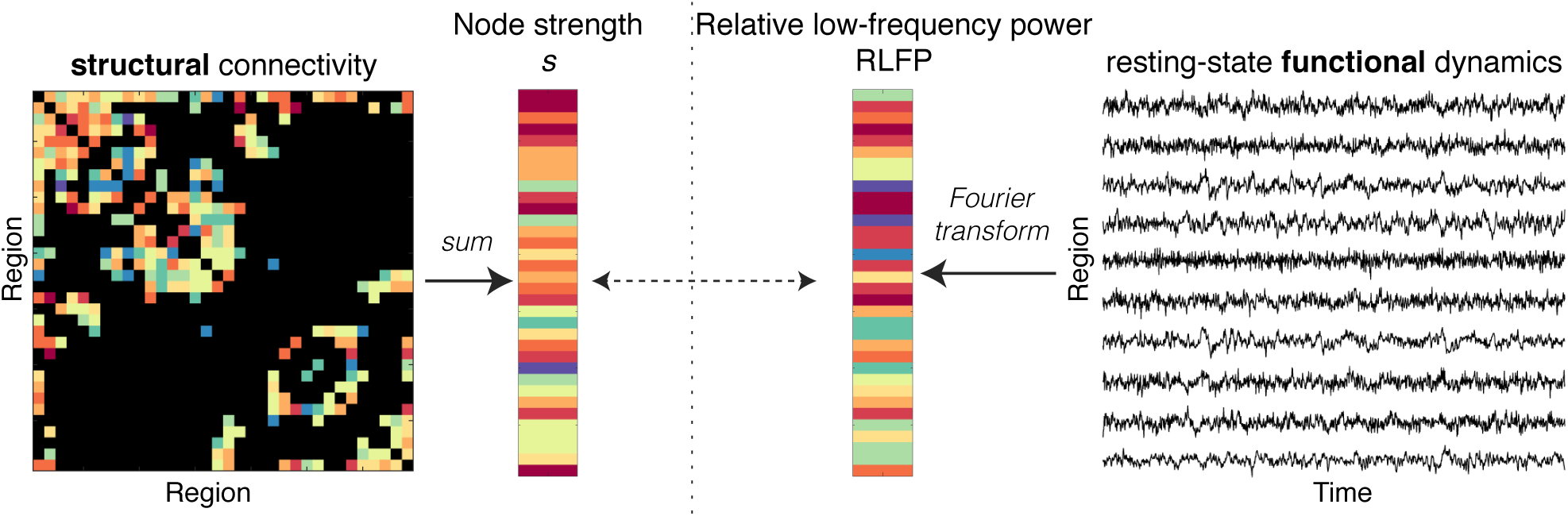
Schematic showing how we investigate the relationship between a region’s structural-connectivity properties to their resting-state dynamics. We summarize each cortical region as its structural connectivity strength, *s*, and its relative low-frequency power, RLFP (*f* < 0.14 Hz). Note that, for the purposes of schematic visualization, edge weights and node strength colors represent relative strength, from low (blue) to high (red).

Based on our previous findings in mouse [47], we summarized BOLD dynamics in a given brain region as the relative low-frequency power, RLFP (*f* < 0.14 Hz). Note that frequencies *f* < 8 × 10^−3^ Hz were removed through high-pass filtering (see Methods). As we use RLFP to understand how the frequencies, or timescales, underlying a given BOLD time series are distributed, we verified that RLFP gives highly correlated results to other common estimates of timescales from time-series data: (i) a fitted exponential decay timescale to the autocorrelation function [24] (Pearson’s *r* = 0.98 across all brain regions, averaged across subjects, *p* = 5 × 10^−26^); and (ii) the area under the autocorrelation function before it passes zero [35] (*r* = 0.995, *p* = 8 × 10^−28^). RLFP is also highly correlated to the similarly constructed metric, fALFF [58, 59] (relative power in the range 0.01 < *f* < 0.08 Hz), which has been widely used to characterize human fMRI (*r* = 0.997, *p* = 2 × 10^−37^). Thus, while we focus our main results on RLFP here, very similar quantitative results were obtained using similar timescale-related statistics.

Relationships between the structural properties of a cortical region and its univariate dynamics were estimated as Spearman correlation coefficients, *ρ*. Region volume, which varies from 49 to 4570 voxels in the Desikan-Killiany Atlas [55], is a major confound, correlating strongly with RLFP, *ρ* = 0.61 (*p* = 2 × 10^−4^; see Fig. S1A), as in the mouse brain [47]. To control for region volume, we computed partial Spearman correlation coefficients, denoted here as *ρ*_*V*_.

### 2.2 Node strength is correlated with power-spectral properties of resting-state BOLD dynamics

We first investigated the group-level relationship between connectivity strength, *s*, and relative low-frequency power, RLFP, by summarizing each brain region as the mean of each quantity across all 100 participants. As shown in Fig. 2A, there is a strong correlation between *s* and RLFP in the left hemisphere of the cortex, *ρ*_*V*_ = 0.53 (controlling for region volume, *p* = 2 × 10^−3^). Similar results were observed in the right hemisphere, *ρ*_*V*_ = 0.57 (*p* = 6 × 10^−4^; see Fig. S3). The positive correlation indicates that human cortical areas with greater aggregate structural connectivity display stronger low-frequency fluctuations, matching the relationship characterized in the mouse brain [47]. Note that RLFP is strongly correlated with region volume, *ρ* = 0.61 (Fig. S1A), and the *s*–RLFP relationship is stronger when region volume is not controlled for, *ρ* = 0.74 (*p* < 2 × 10^−6^; see Fig. S1B).

**Figure 2:**
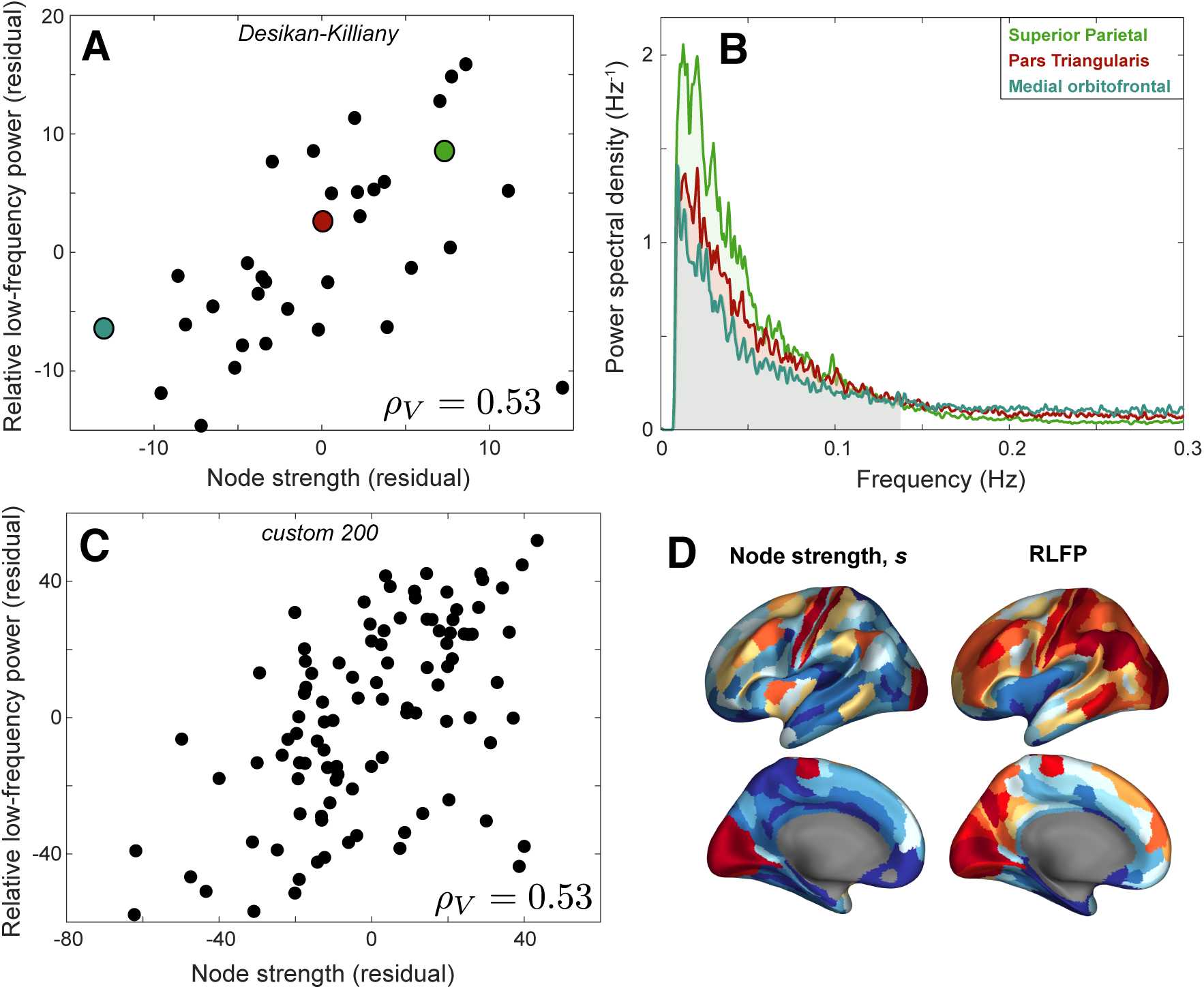
Group-level connectivity strength, *s*, is positively correlated with relative low-frequency power of BOLD dynamics, RLFP (*f* < 0.14 Hz) after correcting for region volume. **A**: Rank residuals of relative low-frequency power (RLFP) and node strength, *s*, across 34 left-hemisphere cortical regions of the Desikan-Killiany Atlas [55], after regressing out region volume. The plot reveals a positive relationship, partial Spearman’s *ρ*_*V*_ = 0.53 (*p* = 2 × 10^−3^). **B** The group-averaged Fourier power spectra for three colored brain areas in A are plotted: medial orbitofrontal area (low *s*, blue), pars triangularis (moderate *s*, red), and superior parietal (high *s*, green), shown up to a maximum of 0.3 Hz. RLFP corresponds to the shaded area under the curve below 0.14 Hz. **C**: As **A**, but for 100 left-hemisphere cortical regions from a custom 200-region parcellation generated by randomly dividing each hemisphere into 100 approximately equal-sized regions [61]. **D**: Spatial maps of node strength and low-frequency power across 180 left-hemisphere cortical areas of the Glasser et al. [62] parcellation, with the relative variation of each metric shown using color, from low (blue) to high (red).

To better understand these findings, we selected three representative brain regions: the medial orbitofrontal region (low *s* = 1.4 × 10^5^), pars triangularis (moderate *s* = 4.6 × 10^5^), and superior parietal cortex (high *s* = 1.7 × 10^6^), as annotated in Fig. 2A. The Fourier power spectrum for each of these brain areas is plotted in Fig. 2B, with the RLFP region shaded (*f* < 0.14 Hz). Differences in spectral power are clearest at low frequencies, especially near the peak power around 0.02 Hz. As the total power is normalized to unity, increased relative power around 0.02 Hz results in lower relative power at higher frequencies. Accordingly, the relationship with *s* is not sensitive to the precise RLFP frequency range, but is reproduced (with opposite sign) at higher frequency bands of the same extent: *ρ*_*V*_ = −0.53 (0.14–0.28 Hz), *ρ*_*V*_ = −0.56 (0.28–0.41 Hz), *ρ*_*V*_ = −0.53 (0.41–0.55 Hz), and *ρ*_*V*_ = −0.53 (0.55–0.69 Hz). The brain region that deviated most from the overall trend was the insula (corresponding to the point in bottom right of Fig. 2A), which has surprisingly low RLFP given its strong structural connectivity across the brain (after correcting for region volume), perhaps due to its vicinity to large blood vessels making it more susceptible to physiological noise [60]. The common statistical summary of univariate BOLD dynamics, fALFF [59], is algorithmically very similar to RLFP and is similarly correlated with *s*: *ρ*_*V*_ = 0.54. Other common measures of timescales derived from the decay of the autocorrelation function (ACF) also yielded similar results, including the decay timescale of Murray et al. [24], *ρ*_*V*_ = 0.48, and the area-based measure of Watanabe et al. [35], *ρ*_*V*_ = 0.51.

The relationship between *s* and RLFP does not depend strongly on cortical parcellation. A similar relationship was found when randomly dividing each hemisphere into approximately 100 equal-sized regions [61], shown in Fig. 2C for the left hemisphere, *ρ*_*V*_ = 0.53 (*p* = 1 × 10^−8^). We also found a significant positive relationship when using the 180-region Glasser et al. [62] parcellation of the left cortex, *ρ*_*V*_ = 0.43 (*p* = 3 × 10^−9^; see Fig. S2), and when resampling the same number of voxels from each brain region (circumventing the need to correct for variation in region volume), *ρ* = 0.43 (*p* = 0.01; see Fig. S1D). Spatial maps of both properties, at the higher spatial resolution of the Glasser et al. [62] parcellation, are shown in Fig. 2D.

### 2.3 Regional structure–function relationships extend across individuals

Having demonstrated a robust group-level relationship between node connectivity strength, *s*, and Fourier spectral characteristics of rs-fMRI time series, we next investigated whether these results could be detected at the level of individual subjects. We measured the partial correlation coefficient, *ρ*_*V*_, between *s* and RLFP for each individual, plotted as a distribution across all 100 individuals in Fig. 3. After false-discovery rate multiple-hypothesis correction [63], 43% of participants individually displayed a significant correlation (*p*_corr_ < 0.05: *ρ*_*V*_ > 0.40). The group-level *s*–RLFP relationship (annotated red in Fig. 2) was stronger than the individual correlation for 91% of participants, consistent with a concentration of meaningful signal (and thus a reduction in measurement noise) through group averaging. To investigate whether inter-individual variation in *s*–RLFP correlation, *ρ*_*V*_, is driven by in-scanner motion, we computed the Pearson correlation between *ρ*_*V*_ and mean framewise displacement across individuals. We found a weak and non-significant relationship, *r* = 0.10 (*p* = 0.3), suggesting that motion is not driving our results.

**Figure 3:**
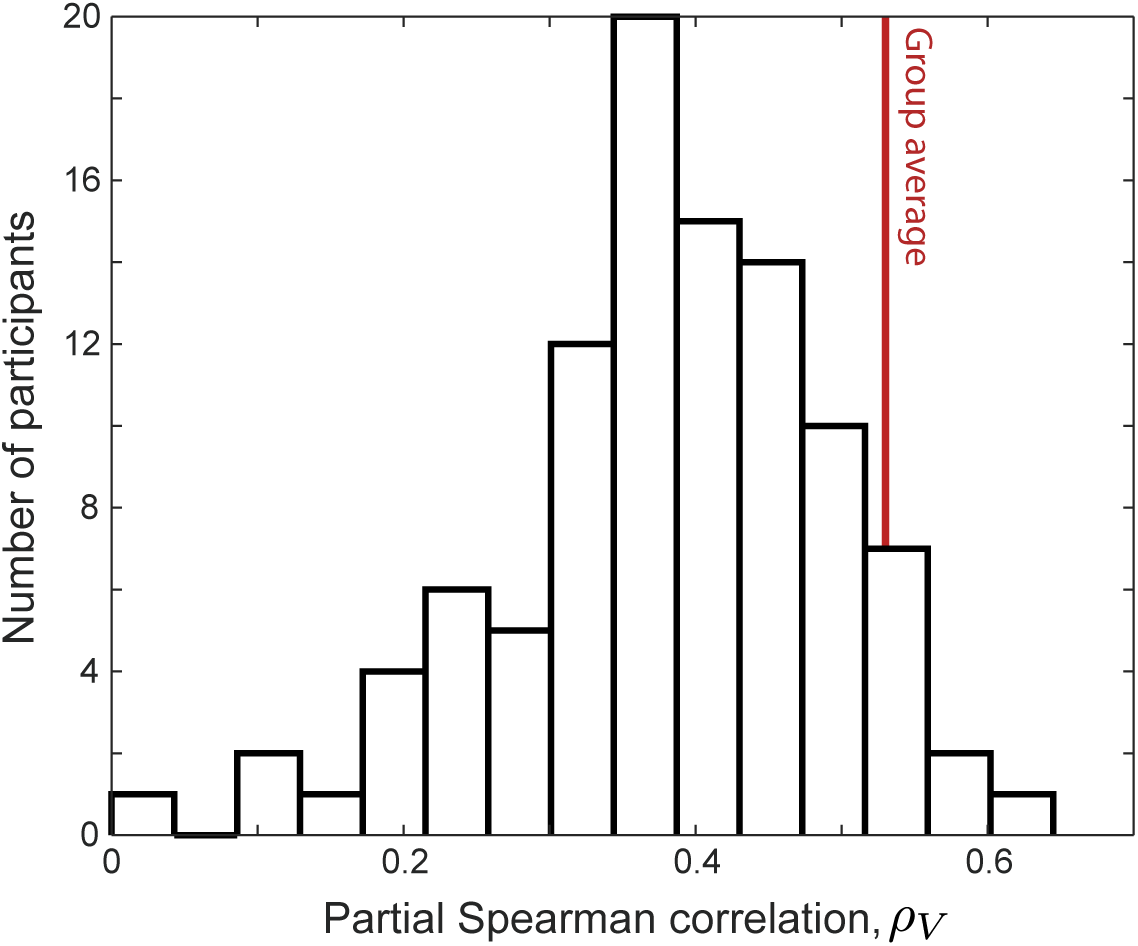
Many individuals exhibit a significant relationship between node strength, *s*, and relative low-frequency power, RLFP. The histogram of partial Spearman correlation coefficients, *ρ*_*V*_, between RLFP and *s* (correcting for variations in region volume) computed separately for each of 100 individuals. The group-level result, *ρ*_*V*_ = 0.54, is shown as a vertical red line.

### 2.4 Diverse properties of BOLD dynamics are informative of node strength

Our results demonstrate that Fourier spectral properties of rs-fMRI, and related measures of autocorrelation timescales [24, 35], are strongly correlated with connectivity strength, *s*. But the time-series analysis literature is vast and interdisciplinary [64]; could other statistical summaries of BOLD time series exhibit stronger relationships to *s*? To investigate this possibility, we used the *hctsa* toolbox [49] to perform a comprehensive data-driven comparison of the performance of 6062 different time-series features. The performance of each feature was measured as *ρ*_*V*_ (computed for each individual and then averaged across individuals). As we are interested in the magnitude of the correlation (not the sign), we took |*ρ*_*V*_ | as the quantity of interest, plotting its distribution across all 6062 time-series features in Fig. 4. While 3768 individual time-series features exhibit a statistically significant partial correlation to *s* (|*ρ*_*V*_ | > 0.39, *p*_corr_ < 0.05), RLFP is amongst those with the highest |*ρ*_*V*_ | (in the top 16% of all *hctsa* features). However, the distribution reveals a tail of alternative time-series features with higher |*ρ*_*V*_ |. Interestingly, these high-performing features recapitulate a familiar set of time-series analysis methods that have previously been used to analyze BOLD dynamics, as well as some unexpected new features. Here we summarize some notable features, labeling them by their name in *hctsa* [49]; for further descriptions of these and other features, see the Supplementary Text (and see Supplementary File 1 for a full list).

**Figure 4:**
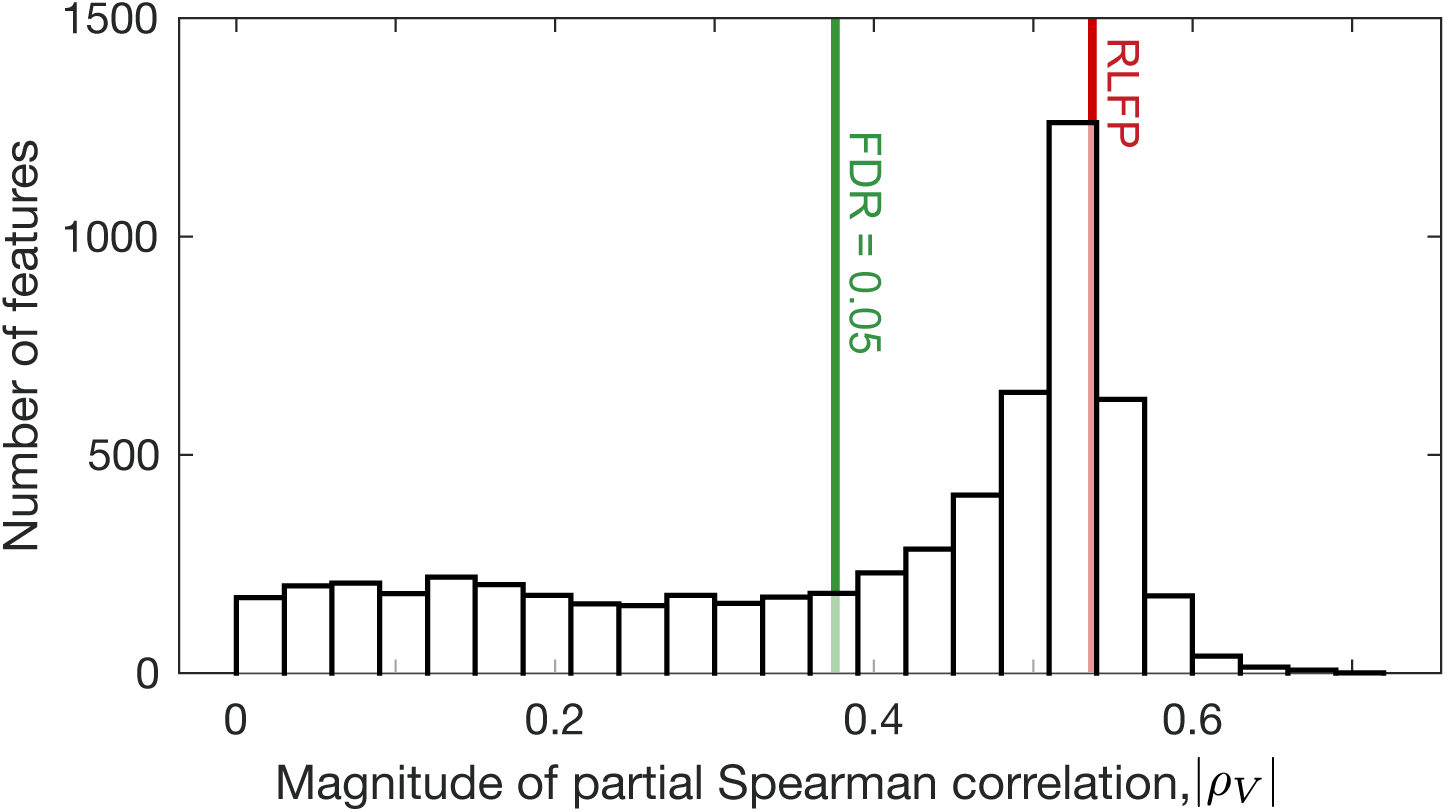
RLFP has amongst the strongest correlations to connectivity strength, *s*, in a comparison to 6062 time-series features. We plot a histogram of absolute partial Spearman correlation coefficients, |*ρ*_*V*_ |, between each of 6062 rs-fMRI time-series features and connectivity strength *s* (controlling for region volume). The features were computed using the *hctsa* toolbox [48, 49]. RLFP (|*ρ*_*V*_ | = 0.53) is shown red, and the 5% FDR-corrected statistical-significance threshold (|*ρ*_*V*_ | > 0.39) is shown green.

Neuroimaging time series have most commonly been summarized as a measure of timescale, derived from the Fourier power spectrum or linear autocorrelation function (which are related via a spectral decomposition, cf. the Wiener–Khinchin theorem) [24, 31, 34, 35, 52]. Our highly comparative time-series analysis highlights many qualitatively similar features. One example is `SP_Summaries_fft_linfitloglog_mf_a2`, *ρ*_*V*_ = −0.59, which estimates the powerlaw exponent of the Fourier power spectrum (as a linear fit in a log-log plot, fitted after excluding the lower and upper quarter of frequencies), reminiscent of Hurst exponent estimation from neural time series [65]. Another interesting high-performing feature is an information-theoretic analogue of the first zero-crossing of the autocorrelation function: the first minimum of the auto-mutual information function [66, 67] computed after differencing the time series: `IN_AutoMutualInfoStats_diff_20_k` *ρ*_*V*_ = −0.63. This incremental differencing step, a common time-series transformation used to stabilize the mean [68], also emerged in a range of symbolic motif features. Symbolic motifs count the frequency of a particular set of consecutive symbols in a time series that has been converted to a symbolic string using a simple coding rule, e.g., coding stepwise increases as ‘U’ (up) and decreases as ‘D’ (down). Motifs associated with small movements are consistent with slow fluctuations and showed positive correlations with *s* (e.g., the ‘AABB’ pattern: `SB_MotifThree_diffquant_aabb`, *ρ*_*V*_ = 0.62), whereas motifs associated with rapid changes exhibited strong negative correlations (e.g., the ‘up-down-up-up’ pattern: `SB_MotifTwo_diff_uduu`, *ρ*_*V*_ = −0.62). The symbolization process may help to capture informative structure in noisy rs-fMRI time series.

Increased durations of rs-fMRI recording have allowed dynamic functional connectivity analyses to characterize changes in functional connectivity across the recording period [69]. Our data-driven analysis highlighted a range of univariate analogues of this concept, flagging a range of high-performing features measuring time-series stationarity. For example, `SY_SlidingWindow_sampen_ent10_2`, *ρ*_*V*_ = −0.65, measures how local estimates of the entropy metric, SampEn(2, 0.1) [70], vary across the time series. *hctsa* also highlighted novel features derived from visibility graphs, which represent each fMRI time point as a node and constructing network edges using visibility rules [71, 72]. For example, *s* is highly correlated to a simple outlier metric of the visibility graph degree distribution, `NW_VisibilityGraph_norm_ol90`, *ρ*_*V*_ = 0.66.

Our results demonstrate the usefulness of *hctsa* in determining the most useful time-series analysis methods for a given problem in an automated, data-driven manner. *hctsa* provides new understanding of how rs-fMRI dynamics relate to structural connectivity strength by flagging an ensemble of time-series features that both encapsulate conventional approaches to analyzing BOLD dynamics, and introducing novel ones.

## 3 Discussion

In this work, we show that the variation of intrinsic fMRI timescales across human cortical areas is related to the variation of structural-connectivity strength, matching a regional structure–function relationship previously observed the mouse brain. This interspecies consistency is observed despite major differences in measurement between mouse (axonal tract tracing and rs-fMRI in 18 anesthetized mice) and human (DWI and rs-fMRI in 100 awake participants). In both species, brain areas with a greater aggregate strength of structural connectivity exhibit slower rs-fMRI BOLD fluctuations, consistent with a hierarchical gradient of intrinsic timescales [21–24]. Our results are robust to cortical parcellation, hold at the level of individuals, and are not driven by motion. We also introduce a highly comparative time-series analysis approach to the problem which, in a data-driven way, recapitulates conventional BOLD signal analysis approaches and highlights a range of promising new temporal features, including symbolic and stationarity-related measures, for characterizing BOLD dynamics. This study expands the investigation of the brain’s structure–function relationship to the level of individual regions (rather than pairs of regions), providing a more complete picture of how the brain’s intricate axonal scaffold shapes spontaneous brain dynamics. Continuing investigations of the structure–function relationship at the level of individual areas will allow us to better understand how the brain’s local circuits shape its dynamics and relate these results to hypotheses about the principles governing the brain’s spatiotemporal organization.

Our results in mouse and human indicate that areas that are more strongly connected to the rest of the brain have slower average timescales. This relationship is consistent with a hierarchy of timescales in which more highly connected areas (high in the hierarchy) serve a more integrative function, integrating diverse information over longer timescales than the fast dynamics and behavioral responses associated with lower sensory areas [22–25, 47, 51, 52]. While many previous studies have reported interareal variation in intrinsic timescales [22, 25, 27, 30–33, 35], to our knowledge, only two prior human studies have related this variation to structural connectivity [31, 34]. Both studies found weak relationships, *r* = 0.12 [34] and *r* = 0.22 [31], using hand-picked dynamical properties of rs-fMRI time series in small sample sizes, and without correcting for regional volume variation. The much stronger correlation reported here, *ρ*_*V*_ = 0.53 (after correcting for region volume), may be due to the high-quality imaging data (1200 volumes at a sampling period of 0.72 s) in a larger sample of 100 individuals. As we discuss later, the low temporal resolution of fMRI is a major limitation of studying timescales, but on these fMRI data, we demonstrate that RLFP, fALFF [59], and measures of the decay of the autocorrelation function [24, 35], are all highly intercorrelated. Given this connection, recent work indicating a significant and strong correlation between timescales estimated from simultaneously recorded EEG and fMRI (*γ* band, adjusted *r*^2^ = 0.71) [35], suggesting that our results may also reflect similar differences at faster timescales. Future multimodal research could probe intrinsic processing timescales to provide a more complete temporal picture of how timescales are structured across the cortex in different species, and its implications for cognition and disease.

While fMRI is most frequently characterized in terms of pairwise correlations (functional connectivity), our results highlight the utility of characterizing local brain dynamics. BOLD dynamics are distinctive to individuals [26], play a role in cognitive function [21], and are disrupted in disease [35]. They are also involved in brain organization, being related to the functional and structural connections of an area, and may provide an indirect measure of its information-processing capabilities [30, 31]. Summarizing the activity dynamics of individual brain areas also yields spatial maps that can be related to other datasets straightforwardly, such as macroscale maps of microstructural variation [38, 73]. While univariate analysis of the BOLD signal is promising, a common problem in analyzing univariate time series is selecting an appropriate analysis method or summary statistic to compute [64]. The neuroimaging literature has most commonly focused on linear autocorrelation properties, either measured directly from the autocorrelation function or Fourier power spectrum. *hctsa* circumvents the need for subjectivity or manual exploration across a vast time-series analysis literature [48, 49], providing a data-driven means of selecting the most relevant types of time-series features for a given problem. By comparing the behavior of thousands of diverse time-series analysis methods, *hctsa* finds interpretable features that vary most strongly with structural connectivity strength, *s*. These features recapitulate conventional timescale-based metrics commonly used in the literature, and also include a range of novel features related to symbolic motifs, stationarity, and visibility graphs. The common theme of applying incremental differencing, a transformation commonly used to stabilize the mean [68], suggests that applying this transformation to BOLD data could enhance the informative signal that can be extracted from it. We also note the prominent performance of time-series stationarity properties, suggesting a fruitful avenue in further characterizing this property, which may loosely be considered a univariate analogue of dynamical functional connectivity [69]. Given the large number of time-series analysis methods suited to long, low-noise recordings, we expect that *hctsa* will be especially useful in discovering informative time-series features from data streams with higher sampling rates than fMRI, such as those measured using ECoG, MEG, and EEG.

This study highlights the ability of direct interspecies comparisons to accumulate evidence for common properties of brain organization. Organizational properties that are shared across scales and species [74] are strong candidates for being under evolutionary selection pressure for serving an important functional advantage. These range from the properties of networks abstracted from the brain’s physical structure—like rich-club connectivity and modularity—through to hierarchical gradients [37, 38] and the patterning of gene expression with structural connectivity [75–77]. The current study adds a regional relationship between structure and function to this set of conserved relationships; future work is needed to establish whether a similar pattern holds across other species, such as *C. elegans* and non-human primates [78], where both structural connectivity and large-scale brain dynamics have been measured.

Relative to rapidly sampled electrophysiological recordings, it can be harder to interpret changes in the power spectrum of a sparsely sampled and noisy fMRI signal in terms of relative timescales. Timescales can be quantified in terms of a natural intrinsic frequency of oscillation, estimated from prominent peaks in the power spectrum in EEG [79], MEG [28], and ECoG [80]. Our coarser estimate of timescale, in terms of power in frequency bands, does not correspond to variations in the power spectrum peak position (see Fig. 2), but rather the relative contribution of low-frequency (relative to high-frequency) fluctuations (*f* ∼ 0.14 Hz) to signal variance. Our analysis of frequency bands across the full power spectrum is motivated by evidence of meaningful neural signal in fMRI at frequencies *f* > 0.1 Hz [81–83]. Given the strong correspondence between RLFP and measures of how the autocorrelation decays with time lag on the fMRI data analyzed here, we are encouraged by recent work demonstrating a close correspondence between autocorrelation-based timescale measurements made in EEG and fMRI [35], suggesting that fMRI signals may be informative of neural dynamics at faster timescales.

Understanding how time-series properties vary across cortical areas has important practical implications for how functional connectivity is estimated and interpreted. We first draw attention to the strong variation of time-series properties with region volume (as more voxel-wise time series are averaged into a region-level time series), and the strong conservation of this relationship between mouse and human [47]. This confound of parcellation is not typically acknowledged or addressed, but is crucial to account for by performing partial correlations using region volume as a regressor, or using parcellations containing regions of equal volume. Our results also suggest that different brain areas may have different intrinsic Fourier spectral properties (and hence distinctive autocorrelation functions). As these differences affect the estimation of functional connectivity, future work should leverage recent statistical developments [53, 84, 85] to better account for regional variations in BOLD autocorrelation when estimating and performing inference on functional connectivity.

We quantified connection weights in our connectomes using streamline count for consistency with many other studies in the human connectomics literature. This measure is conceptually closer to the normalized connection weight measure used in our prior study of the mouse [47]. It is well-known that streamline count and other tensor-based metrics, such as mean tract fractional anisotropy, are only indirect proxies for the actual number (or integrity) of axons connecting two regions [86]. Exploring how regional dynamics relate to other properties of axonal connectivity measured with more biologically informative metrics would be a fruitful avenue for future work. While fMRI data-processing pipelines have been shown to have a large impact on resulting functional connectivity estimates [87], it will be important for future work to take similar care in understanding how preprocessing steps affect the univariate properties of fMRI data.

Our findings suggest that the brain may exploit timescales to efficiently store, process, and transfer information, with these timescales coupled to the underlying structural connectivity properties of a brain area [30] in the same way in the mouse brain and human cortex. An intriguing possibility is that the relationship reported here reflects a causal role of long-range structural connectivity in shaping the intrinsic activity timescales of a brain area, as has been demonstrated in simulations of network-coupled dynamical systems [32, 51, 52]. This is consistent, for example, with greater aggregate inputs to a brain region pushing it towards its critical point where dynamical timescales are slower [88]. Our results could also be explained by a hierarchical gradient of microstructural variation [37, 38], along which both structural connectivity strength and spontaneous dynamics vary [19, 27, 28]. In this view, timescales vary hierarchically (due to variations in cortical microstructure or subcortical inputs), but may not be causally modulated by corticocortical structural connectivity strength. Regional variations in perfusion, perhaps to support the increased metabolic demands of highly-connected hub areas [89], could also manifest in corresponding spatial differences in BOLD dynamics [60, 90]. In the absence of experiments that can causally manipulate structural connectivity, computational modeling will continue to play a crucial role in distinguishing between possible mechanistic explanations of the statistical relationships characterized here, towards an understanding of the general and specific mechanisms through which intrinsic timescales may be shaped. Given the cognitive importance of how differences in local neural dynamics, including their timescales, are organized across the cortex [33, 35], understanding the physical mechanisms shaping variations in BOLD dynamics could lead to novel new treatments that aim to rectify abnormal timescales in the brain, e.g., using a transcranial magnetic stimulation (TMS) protocol [42] tailored to an individual’s structural-connectivity profile. Integrating data across species and scales to elucidate common relationships, and using theoretical modeling approaches to propose possible mechanisms underlying those patterns, will be key to understanding how the brain’s organization allows it to efficiently process and integrate information.

## 4 Data and Methods

Code for reproducing our analyses is at https://github.com/johnfallon/humanStructureFunction. Data to support the findings of this study are available from the HCP at https://db.humanconnectome.org.

### 4.1 Data acquisition and preprocessing

MRI data were downloaded from the Human Connectome Project (HCP) [54]. We selected the HCP 100 unrelated participants dataset (54 males, 46 females) for detailed analysis [54], as in previous work [91]. All participants were healthy and aged between 22–35 years and provided written informed consent; ethics was approved by the Institutional Review Board of Washington University. We used the minimally preprocessed data of which full details can be found elsewhere [92]; a broad overview is provided here.

#### 4.1.1 Diffusion-weighted imaging

A 3T Siemens Skyra scanner with a customized head coil (100 mT/m maximum gradient strength and a 32 channel head coil) located at Washington University, St Louis, was used to acquire all neuroimaging data. Diffusion data were acquired using a spin-echo EPI sequence with the following parameters: TR/TE = 5520/89.5 ms, slice thickness = 1.25 mm, 111 slices, 1.25 mm isotropic voxels. Three gradient shells of 90 diffusion-weighted directions and six b0 images were collected with right-to-left and left-to-right phase encoding polarities for each of the three diffusion weightings (1000, 2000, and 3000 s/mm^2^). For additional imaging parameters see Glasser et al. [92]. The diffusion data had been pre-processed using the HCP diffusion pipeline [92], which included normalization of b0 image intensity across runs, correction for EPI susceptibility and eddy-current-induced distortions, gradient-nonlinearities, subject motion and application of a brain mask.

Subsequent processing of the diffusion data used `MRtrix3` [56] and FMRIB Software Library [57]. Tractography was conducted using Fibre Assignment by Continuous Tracking (FACT), a deterministic measure [93, 94] This deterministic measure was selected over probabilistic methods because it is less prone to false-positive connections [95], which have been shown to be more detrimental to network construction than false negative connections [96]. A total of 10 million streamlines were generated with a step size of 0.125 mm. Streamlines terminated when the curvature exceed 45°, when the fractional anisotropy value was less than 0.1, or if the length was greater than 250 mm.

In order to further improve the biological accuracy of the structural networks, Anatomically Constrained Tractography (ACT) and Spherically Informed Filtering of Tractograms (SIFT-2) were applied alongside and to the tractography data. ACT delineates the brain into different tissue types (e.g., cortical grey matter, subcortical grey matter, white matter, CSF). This information is then used while tractography is being conducted to ensure streamlines are beginning, traversing, and terminating in anatomically correct locations [97]. Another issue hampering tractography is the density of reconstructed connections is not reflective of the underlying diffusion data [98]. SIFT-2 addresses this limitation by modeling the expected density of connections as calculated from the diffusion signal before comparing this prediction to the connection densities obtained in tractography. Streamlines are then weighted by a cross-sectional area multiplier determined by this model fit [99]. This same model of diffusion density was also used to dynamically choose streamline seeding points during tractography [99].

Our main cortical parcellation was the 68-region Desikan–Killiany Atlas [55] (34 regions per hemisphere). To demonstrate robustness of our results, we also compared two additional parcellations: (i) Glasser et al.’s 360-region HCP parcellation (180 regions per hemisphere) [62], and (ii) a custom built 200-node parcellation (100 regions per hemisphere) which was formed by randomly dividing each hemisphere into 100 approximately equal-sized cortical regions [61]. These parcellations were generated on the Freesurfer-extracted surfaces for each subject and then projected to volumetric space.

As in the mouse [47], we focused our analysis on a single hemisphere. Analyzing ipsilateral connectivity also has the advantage of avoiding errors associated with reconstructing long-range contralateral connections using diffusion tractography [Reveley et al., 2015]. Ipsilateral structural connectivity within the left hemisphere was represented as a weighted, undirected 34 × 34 adjacency matrix, *A*_*ij*_, where each entry captures the number of streamlines with termination points within 5 mm of either regions *i* and *j*. A group-weighted structural connectome, *G*_*ij*_, was constructed by retaining inter-regional connections that were present in more than 75% of participants [100], and setting edge weights to the average value across participants (where zero entries were not included in the average). The resulting adjacency matrix had an edge density of 25%, and edge weights varying from 322 to 7.6 × 10^4^. Following Sethi et al. [47], each brain region was summarized as its connectivity strength, *s*, calculated by summing all streamlines connected to a region (after applying ACT and SIFT-2).

#### 4.1.2 Resting-state fMRI

Resting-state fMRI (rs-fMRI) data were downloaded from the HCP database [54]. Images were obtained using a gradient-echo, echo planar image (EPI) sequence with the following parameters: TR/TE = 720/33.1 ms, slice thickness = 2.0 mm, 72 slices, 2.0 mm isotropic voxels, frames per run = 1200. We used the volumetric EPI data from the first rs-fMRI session (left-right phase encoding), processed and denoised using ICA-FIX [101].

Subsequent processing of the rs-fMRI data was performed. First, the rs-fMRI time series were linearly detrended. Then, to provide more stringent control over nuisance signals we regressed the rs-fMRI data against mean white matter (WM) and mean cerebrospinal fluid (CSF) as well as the global signal (GS) using `fsl_regfilt`. Specifically, grey matter (GM), WM, and CSF probability masks were generated using SPM8’s New Segment routine. The WM and CSF masks were thresholded and binarized, retaining only voxels with > 99% probability. The GM mask was thresholded and binarized to retain only voxels with > 50% probability. The binary GM mask was then subtracted from the binary WM and CSF masks to ensure no gray matter voxels were present in the estimation of the WM and CSF nuisance signals. Estimating the GS was done by taking the union of two whole brain masks created using FSL’s `bet` function applied to the spatially normalized EPI and T1-weighted images [87]. All nuisance time series were extracted by taking the mean over all voxels in the respective masks. Finally, we removed low-frequency (*f* < 8 × 10^−3^ Hz) fluctuations using a high-pass filter as a hard threshold at 8 × 10^−3^ Hz, applied to the EPI data via a fast Fourier transform. Once processed, EPI time series were summarized at the level of brain regions by averaging voxel time series over all voxels within each parcel.

### 4.2 BOLD time-series analysis

For each brain region (34 left-hemisphere regions for our default parcellation) in each subject, we extracted a BOLD time series. Following Sethi et al. [47], we focused our main analysis on the power across frequency bands of the discrete Fourier transform of each BOLD time series. BOLD time series were linearly detrended and normalized to unit variance using a *z*-score before applying a fast Fourier transform. Variance normalization ensures that the total power in the Fourier power spectrum is unity; the power in a given frequency band represents relative power and is unitless. In this work, we refer to relative low-frequency power (RLFP) as the proportion of power contained in the lowest 20% of frequencies (*f* < 0.14 Hz), as in Sethi et al. [47].

For initial comparison to some commonly used time-series metrics, we took an implementation of fALFF based on the REST toolkit [102] (see `SP_fALFF` in the code repository), and our implementations of the autocorrelation function decay timescale and area are taken from `CO_AutoCorrShape` from the *hctsa* toolbox, available at https://github.com/benfulcher/hctsa [48, 49]. To more comprehensively investigate how the power in specific frequency bands of the Fourier power spectrum compares to alternative univariate time-series properties, we compared across the full set of time-series features in *hctsa* (v0.96) [49]. This software was used to extract over 7000 features from each rs-fMRI time series. Following standard procedures [49], we filtered features that returned special values (features inappropriate for these data) or were approximately constant across all brain regions (features provide no meaningful information). We then restricted our analysis to 6062 well-behaved features that were not filtered from any participant.

To investigate the dependence of our results to variations in region volume [47], we estimated the volume of each region in our parcellation by summing the number of 0.7 mm isotropic voxels in a region using the T1-weighted image. Region volume was controlled for by computing a partial Spearman correlation. We used Spearman rank correlations due to the frequently non-normally distributed nodal properties, particularly region volume and node strength.

## Acknowledgments

The authors would like to thank Leonardo Gollo and Dan Lurie for thoughtful comments on a draft manuscript. AF was supported by the Sylvia and Charles Viertel Foundation. BDF was supported by an NHMRC Fellowship (1089718).

## Notes

### Competing Interest Statement

The authors have declared no competing interest.

