## Supplementary Text for "Timescales of spontaneous fMRI fluctuations relate to structural connectivity in the brain"

### Supplementary Results

#### Controlling for region volume

The cortical parcellation used here displays large variation in region size, from 49–4570 voxels. As well as our main analyses, where this variation is controlled for as a partial Spearman correlation, $\rho_V$, we conducted an additional analysis that randomly sampled 49 voxels from every region, circumventing the need to correct for volume. Even after this dramatic loss in data, we still measured a significant correlation between node strength, $s$, and relative low-frequency power, RLFP, $\rho = 0.43$ ($p < 1 \times 10^{-3}$).

#### Diverse rs-fMRI time-series features are informative of connectivity strength

We used a data-driven method, *hctsa* [1, 2], to compare the performance of a comprehensive library of time-series analysis methods, and thereby contextualize the performance of RLFP. As well as highlighting the usefulness of time-series features derived from the linear autocorrelation function and power-spectrum to analyze neural timescales, the comparison highlighted a range of novel metrics related to these properties and others. All features are listed in Supplementary File 1. Some notable examples are summarized below:

**Stationarity** Measures of stationarity (loosely: how the time-series properties change across the recording period) featured heavily amongst the most informative time-series properties of connectivity strength, $s$. High-performing features captured this in different ways, including: (i) how the standard deviation predicted by time-series models reproduced the standard deviation in the real data across subsegments of the time series ($\rho_V = -0.69$ to $\rho_V = -0.67$); and (ii) `SY_SlidingWindow_sampen_ent10_2`, $\rho_V = -0.65$, which measures how local estimates of $\mathrm{SampEn}(2, 0.1)$ vary across the time series.

The flagging of stationarity features by *hctsa* is interesting given that these types of properties may only be measurable in long time series like those analyzed here: HCP rs-fMRI data are unique in their length (1200 samples) and relatively high sampling rate (TR = 720 ms).

**Fourier power spectrum** Classical features related to properties of the autocorrelation function and the Fourier power spectrum were flagged by our analysis. For example, `SP_Summaries_fft_linfitloglog_mf_a2`, $\rho_V = -0.59$, measures the gradient of the Fourier power spectrum represented as a log-log plot (fitted excluding the lower and upper quarter of frequencies).

**Symbolic motifs** Symbolic motifs symbolize a time series, e.g., coding stepwise increases as 'U' (up) and decreases as 'D' (down), turns a time series into a string of letters (e.g., 'UUDUDDDUD...'). Measuring the proportion of different motifs in this string provides a simple, intuitive, and noise-robust way of capturing recurring patterns, with repeats of words like 'UUUU' and 'DDDD' reflecting a slow-varying process, and words like 'UDUD' reflecting a fast-varying process. Node strength was highly correlated to features like `SB_MotifThree_diffquant_bbcc`, $\rho_V = 0.66$: the frequency of the 'bbcc' motif in a difference-based symbolization of the data into an equiprobable three-letter alphabet (with the largest negative deviations coded as 'a', moderate deviations coded as 'b' and large positive deviations coded as 'c'). Similar results were obtained for conceptually similar features, with positive correlations for symbolic motifs representing slow fluctuations, e.g., `SB_MotifThree_diffquant_aabb`, $\rho_V = 0.62$, and negative correlations for symbolic motifs representing faster fluctuations, e.g., `SB_MotifTwo_diff_uduu`, $\rho_V = -0.62$ (the 'up-down-up-up' motif in a two-letter alphabet).

**Visibility graph** Visibility graphs convert a time series into a network, representing each time point as a node, and constructing edges using visibility rules [3]. Connectivity strength, $s$, was highly correlated to a simple measure of outliers in the degree distribution of visibility graphs extracted from the fMRI time series [3, 4], `NW_VisibilityGraph_norm_ol90`, $\rho_V = 0.66$. Many other properties of the visibility graph (and horizontal visibility graph) were predictive of connectivity strength, $s$, including extreme value fits to the degree distribution (e.g., `NW_VisibilityGraph_horiz_evparm2`, $\rho_V = -0.58$) and goodness of fit of a Gaussian distribution (`NW_VisibilityGraph_horiz_gaussnlogL`, $\rho_V = -0.58$) and the standard deviation of the degree distribution (`NW_VisibilityGraph_horiz_stdk`, $\rho_V = -0.58$).

### Supplementary Figures

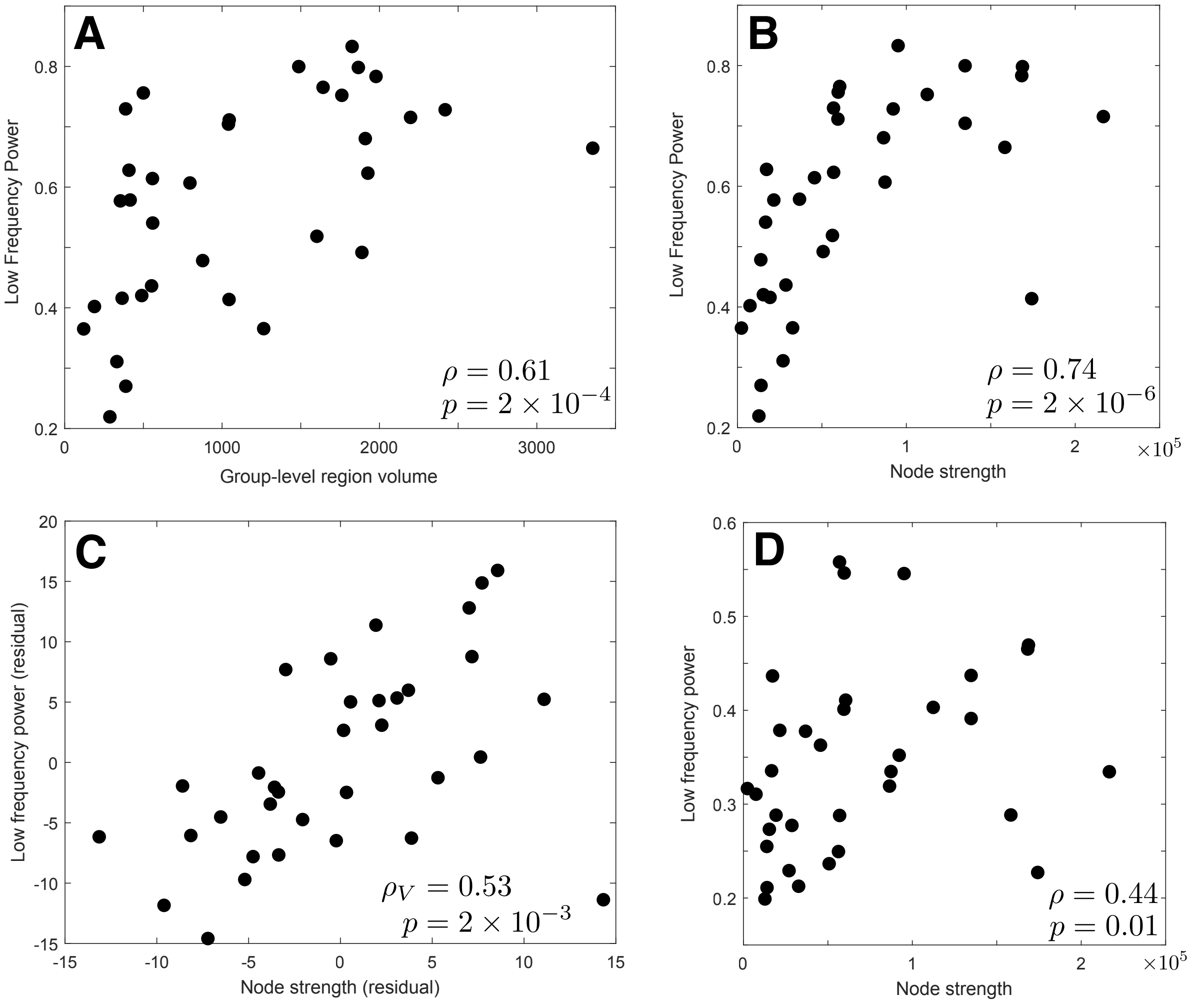


Figure S1: **Relative low-frequency power (RLFP) is strongly correlated to region volume.** Scatter plots of RLFP as a function of: **A** region volume, **B** Node strength, $s$, **C** Node strength rank residuals (controlling for region volume), and **D** Node strength using a parcellation in which each region's time-series are averaged over 49 randomly selected voxels.

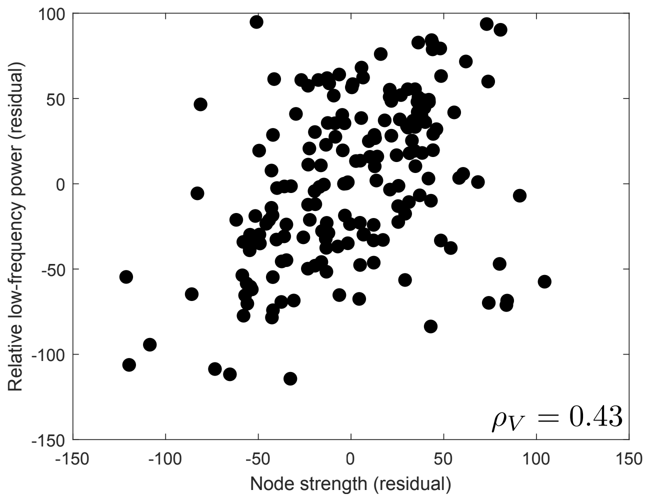


Figure S2: **The relationship between relative low-frequency power (RLFP) and node strength, $s$ holds in the Glasser et al. [5] parcellation.** We plot a scatter plot of the rank residuals of RLFP and $s$ in the left hemisphere using a 360-region parcellation [5] (180 regions in left hemisphere): $\rho_V = 0.43$, $p = 3 \times 10^{-9}$.

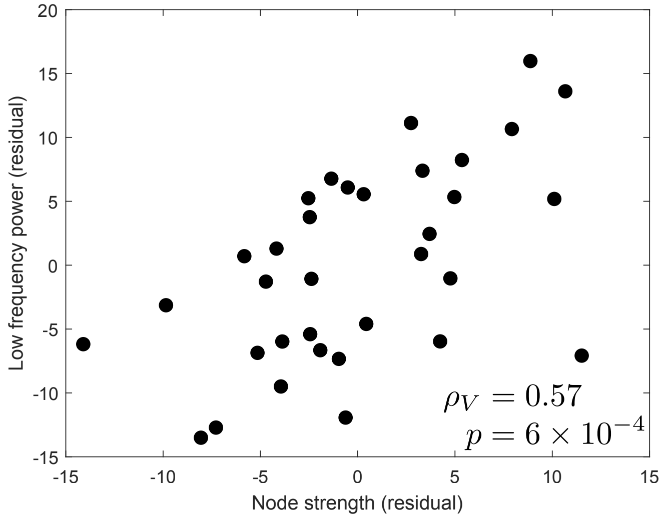


Figure S3: **The relationship between node strength, $s$ and relative low-frequency power, RLFP, holds similarly in both the left and right hemispheres.** Here we regenerate plot of Fig. 2A in the right hemisphere, showing rank residuals of relative low-frequency power (RLFP) and node strength, $s$, across 34 right-hemisphere cortical regions of the Desikan-Killiany Atlas [6], after regressing out region volume. The plot reveals a positive relationship, partial Spearman's $\rho_V = 0.57$ ($p = 6 \times 10^{-4}$).
